# Exact graph-based analysis of scientific articles on clinical trials

**DOI:** 10.1101/164475

**Authors:** Ilya Levin

## Abstract

This article describes Amorpha, a software package based on new concept of exact graph-based linguistic analysis. Analytical capabilities of Amorpha are demonstrated using analysis of scientific abstracts on clinical trials from PubMed. Current trends in therapy of breast cancer and psoriatic arthritis were analyzed using 400 abstracts on breast cancer and 131 abstracts on psoriatic arthritis. The spectrum of diseases that currently treated with paclitaxel was extracted from 400 most recent abstracts on paclitaxel.

In addition to text representation, analytical results are presented as graph images showing essential concepts of a text. Amorpha is not designed specifically to analyze clinical trials and will be also useful for analysis of biological scientific articles and regulatory documents. Amorpha does not require any preliminary knowledge base, ensures full coverage of target text and 100% accuracy of obtained results.

## Introduction

The procedures of information analysis are usually performed within few seconds in relational database and easily expressed in SQL language. However, in real work with clinical scientific and regulatory documents essentially the same procedures are performed within few hours or few days. This work is usually aimed to find some information on selected issues in multiple documents (may be using complex criteria), compare and analyze relevant information, and draw conclusions.

PubMed offers only the tools for search; the analysis of scientific articles and other relevant literature is still performed manually and requires significant time and efforts. On the other hand, widely accepted methods of linguistic analysis are based on probabilistic algorithms. The use of probabilistic methods is explained by obvious difficulties in computational analysis of natural language. These difficulties include such well known phenomena as ambiguity (one word can correspond to different meanings) and variance (several words for one meaning). In addition, there are very significant differences in writing style. In particular, some ideas can be expressed implicitly. This makes computational analysis even more complex. Finally, a text can contain mistakes.

When a scientist works with biological or medical articles, he needs to retrieve essential relevant information, buried somewhere in these articles. Probabilistic tools for text analysis **<u>by definition</u>** will not produce exact result. And this huge amount of work is still performed manually.

This article describes Amorpha, a software package based on a new concept of **<u>exact</u>** graph-based analysis. The use of Amorpha software tools for analysis of a big text or a collection of texts (corpus) dramatically accelerates and facilitates the process of information retrieval. For example, a corpus, comprised of multiple source documents, corresponding to total 100 - 300 pages, can be evaluated for key concepts typically within some few minutes. The system provides representation of the issues that actually exist in a corpus. The corpus can be examined using preliminary defined terms, as well as with new terms, identified by Amorpha software as being of key importance for evaluated corpus. Moreover, Amorpha allows to find new important trends.

Analytical capabilities of Amorpha are demonstrated using analysis of scientific abstracts on clinical trials. The abstracts were obtained from PubMed, and essential information was retrieved. In addition to text representation, this article demonstrates the analytical results as graph images showing essential concepts of a text in easy and intuitive form. There is no need for any preliminary knowledge base: a text is analyzed just as it is. The system is not designed specifically to analyze clinical trials and will be also useful for analysis of biological scientific articles and regulatory documents.

Before publication of this article, Amorpha core technology was validated for 7 years in wide range of texts. The system is always helpful, regardless of language, content and complexity of a text. Amorpha does not require on any preliminary knowledge base, ensures full coverage of target text and 100% accuracy of obtained results.

### Software description

Amorpha reads a text and splits it into <u>singles</u>, elementary units of a text. These singles include words, digits, numbers, percents, and non-word symbols, such as punctuation signs, brackets, quotes, etc. Then the program produces a list of unique <u>reference singles (**Refsi List**)</u>, ordered by the frequency of these singles in current text, in descending manner (Picture 1). Therefore, the most important words are always in the top of Refsi List.

**Picture 1.**
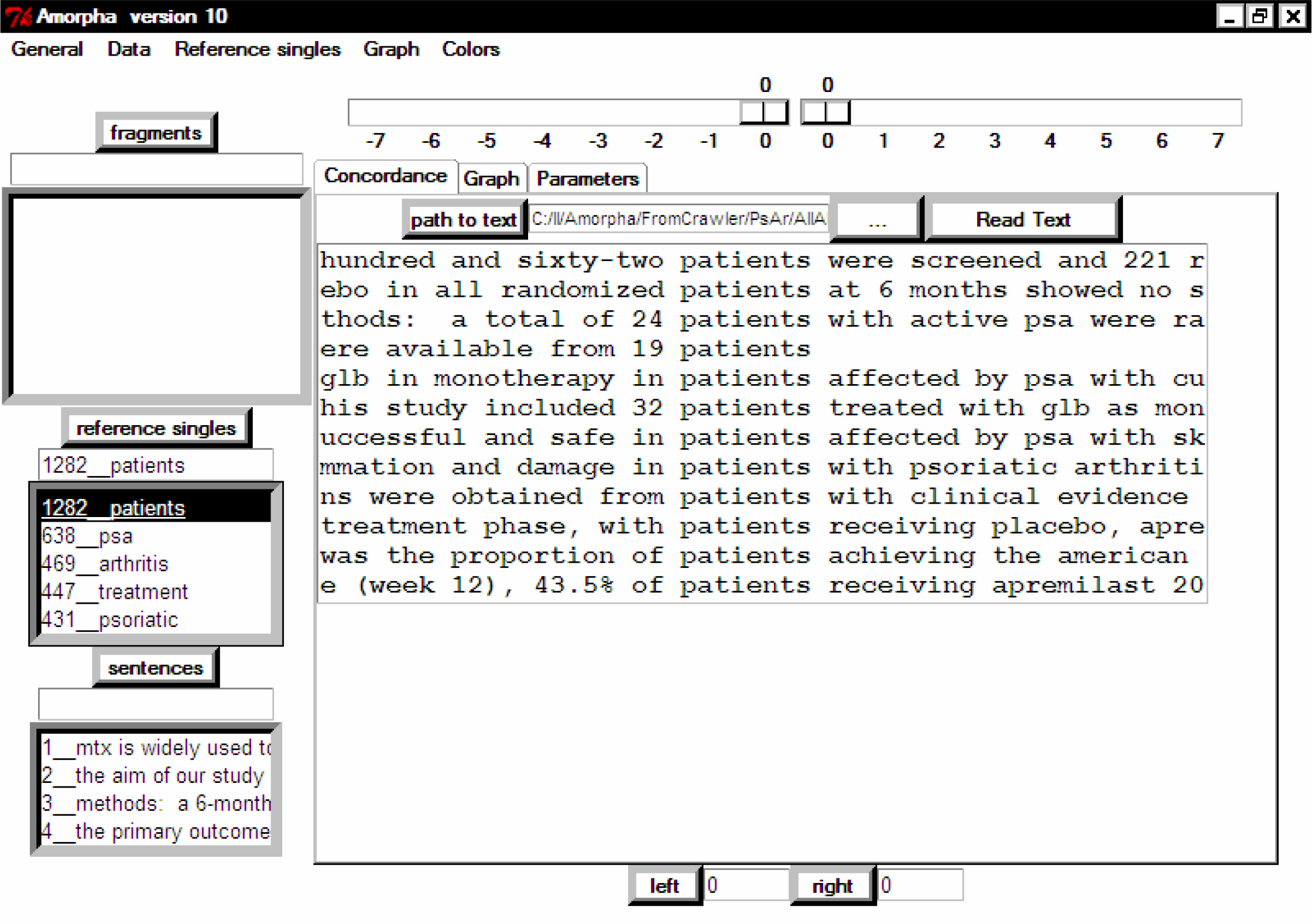
Amorpha version 10 shows Reference Singles list with concordance.

The following singles were excluded from the list of reference singles:

- punctuation signs, brackets and other non-word symbols (except digits, numbers, and percents)
- conjunctions, prepositions, articles
- modal verbs (in some cases)

These singles were excluded from Refsi List because usually they have high frequency (“heavyweight”) and attract our attention on the graph, while not reflect essential concepts of analyzed text. All digits, numbers, and percents remain in the text, because these singles may be important for understanding of this text. All singles are switched to lower register (no capital letters allowed) to ensure recognition of identical singles.

Each reference single is connected to a list of corresponding sentences. For example, the single “study” is connected to the list of all sentences that contain the word “study”. Moving down on the Refsi List from the top provides the context of each currently selected single (word, digit, percent, abbreviation, etc.) in <u>all sentences</u> of current text in one window. Thus, instead of walking on separate parts of texts, we can see all relevant context together. This context, centered by selected single, is known in linguistics as <u>concordance</u>.

Representation of a text with Refsi List and concordance is shown in the picture below. <u>PsAr Corpus</u> (collection of 131 abstracts on psoriatic arthritis) was used as the source text.

Walk on Refsi List from the top down provides the context of selected singles **<u>according to their importance.</u>** This is the key moment in Amorpha programs. Therefore, a scientist, working with the program, can focus his attention primarily on the important (“heavy-weight”) words. In fact, frequency distribution of words **<u>always</u>** has hyperbolic shape and follows power law. This principle is known as Zipf’s law [1] of word distribution. In economics, this distribution is described as “Paretto distribution” (80-20 principle).

Pictures below (Picture 2 and Picture 3) demonstrate the shape of word frequency distribution curve, obtained for PsAr Corpus.

**Picture 2.**
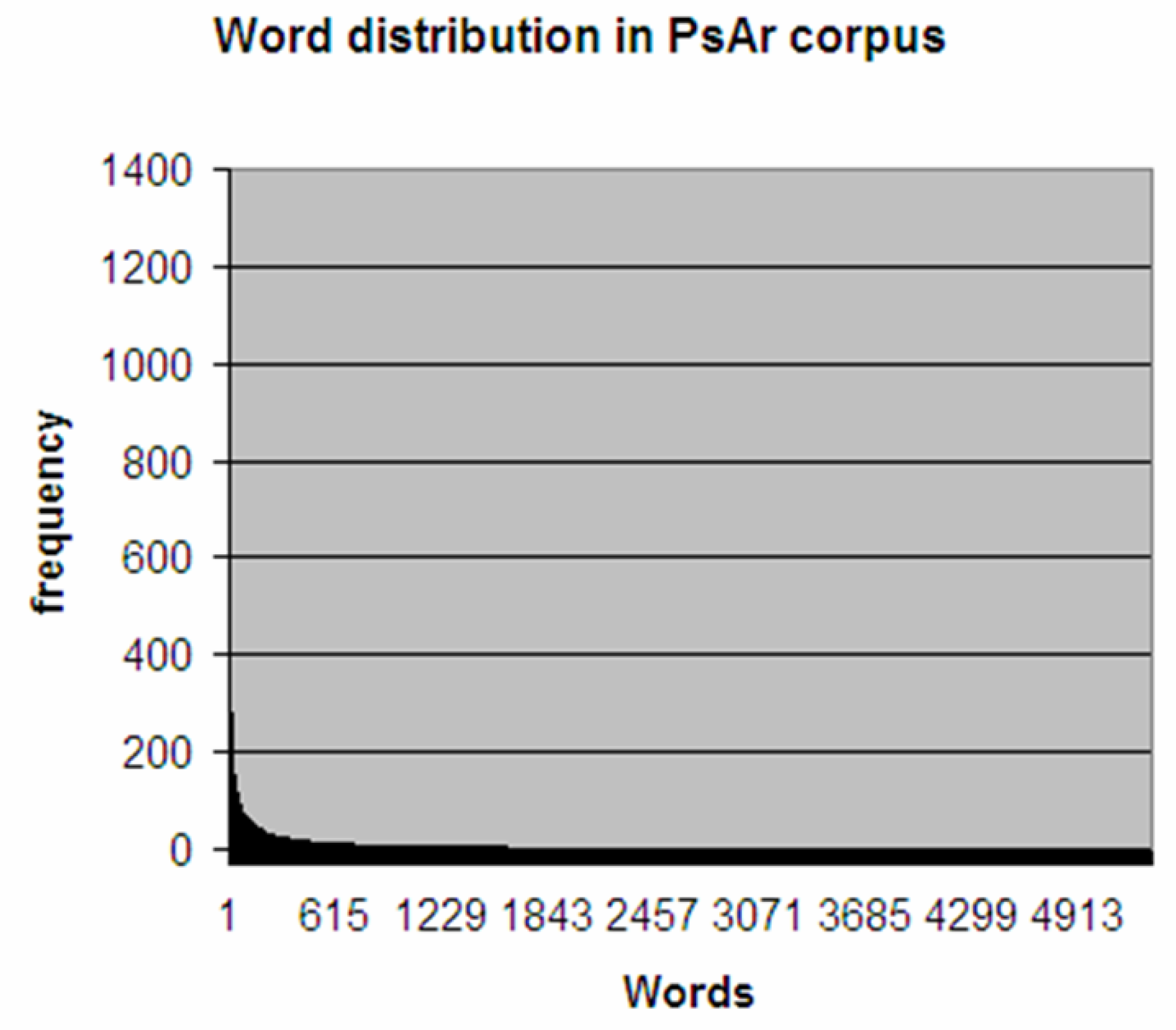
Words distribution of PsAr corpus. This picture shows long tail of low-frequency singles

**Picture 3.**
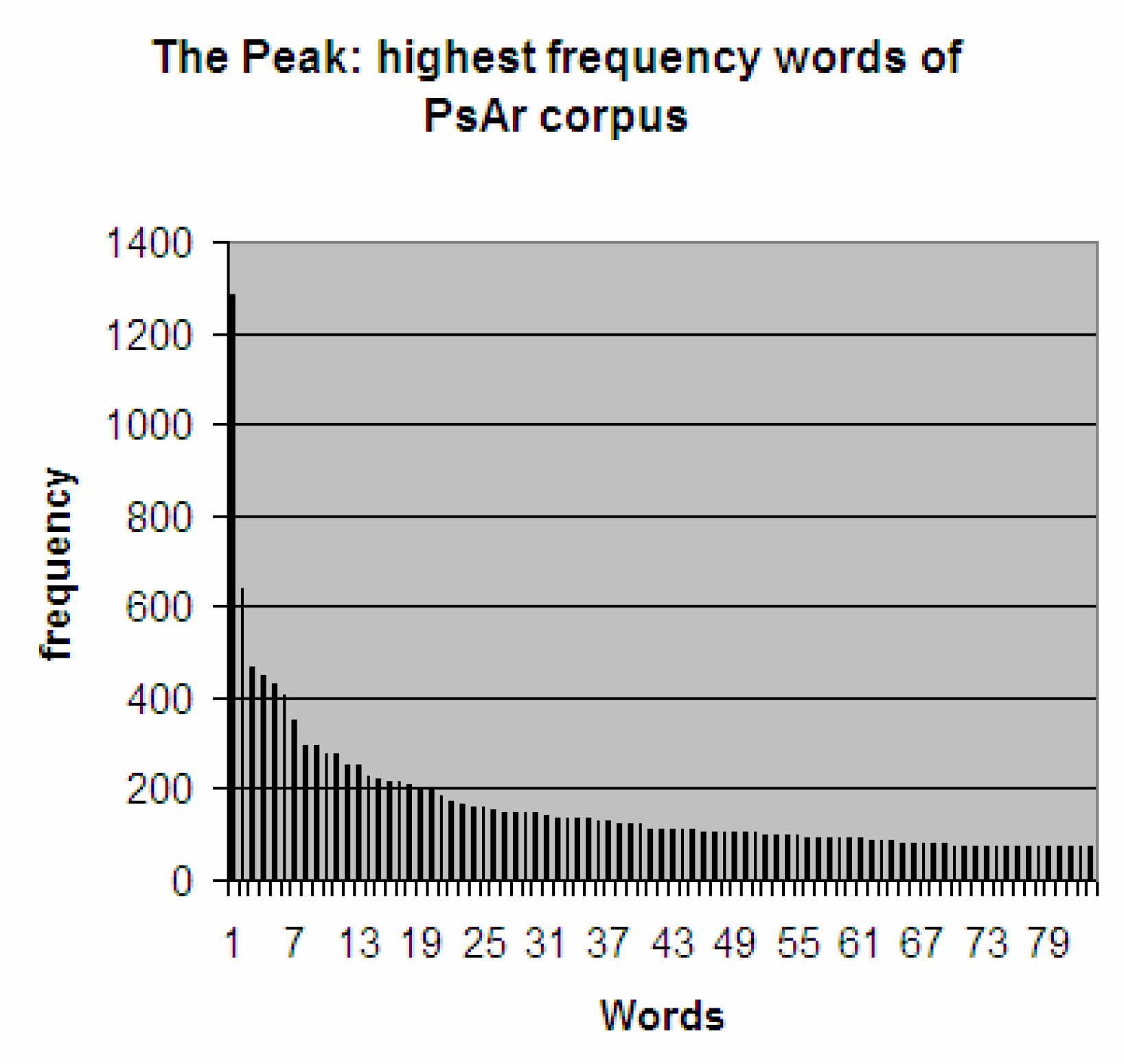
The peak: highest frequency words: minimal word frequency is limited to 71. The area near Y axis is occupied by singles with very high frequency. Top 20 of highest frequency singles is shown in Table 1.

Hyperbolic distribution curve can be divided to 3 components. The first components is high and narrow head (the <u>peak</u>) that approximates (almost in parallel to) the Y-axis. The second component is the <u>middle</u> (continuous curve). The third component is long <u>tail</u> that approximates the X-axis.

The <u>peak</u> always contains the most important words of current text. Usually, these words are not ambiguous and have one meaning in the text. These top words represent the most important terms of the text (Table 1)\.

**Table 1.**
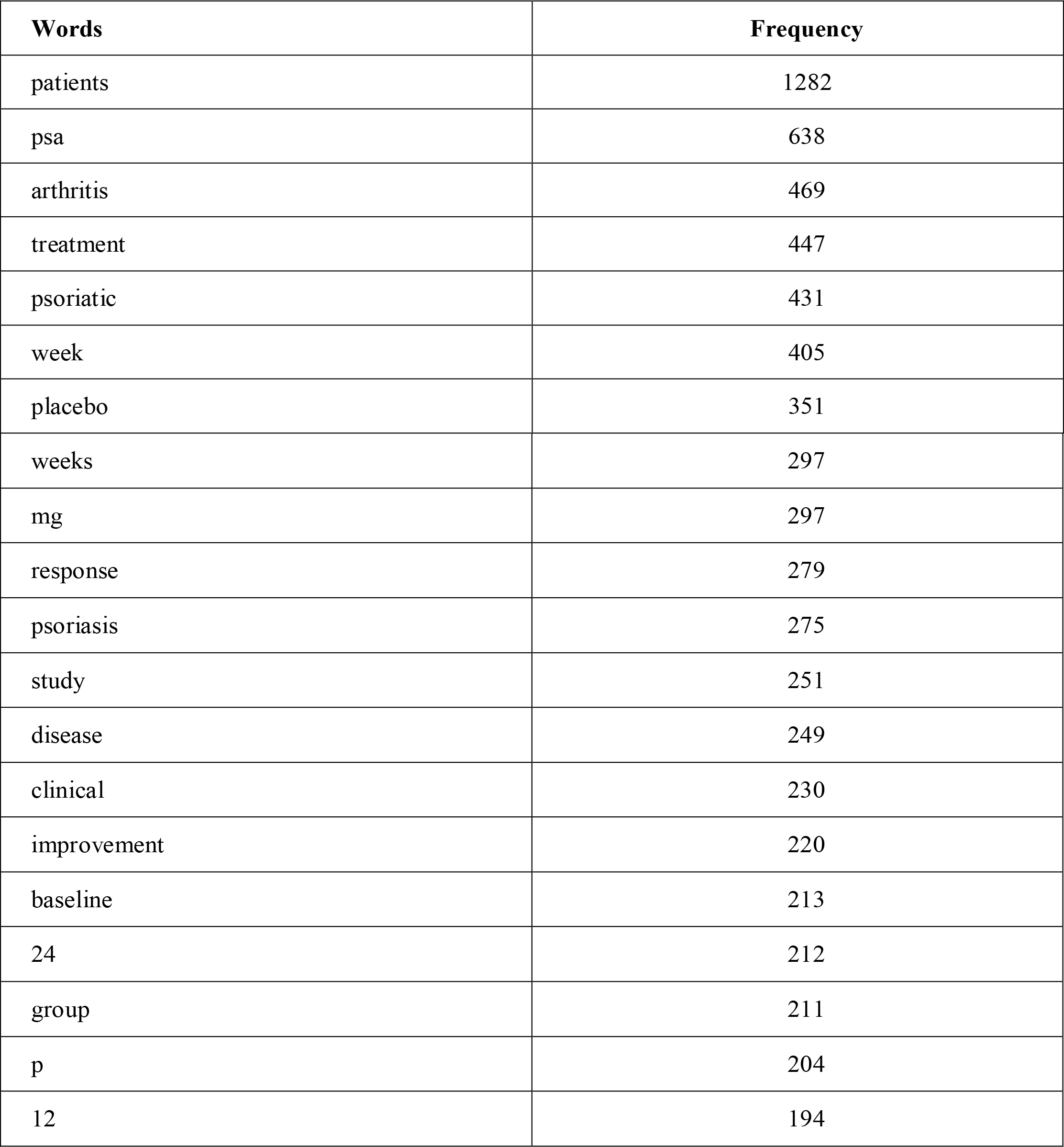
Top 20 singles of PsAr corpus

The <u>middle</u> part of distribution curve contains words with moderate frequency. The low-frequency (from 3 to 1) words and non-word singles are located in the <u>tail</u>. The tail always seems to be very long. Actually, these words consist only about 10% of total amount of words (without any normalization by frequency). Therefore, examination of peak and middle regions <u>ensures</u> control on the majority of words in **<u>any</u>** text.

The walk on frequency-sorted list of words with examination of concordance is actually equivalent to the walk on a big <u>summary graph</u>, obtained directly form the text. This idea will be explained in the following sections.

## Graph-based analysis of big networks

During the past years, the graph-based methods, developed for social network analysis, have been used to model different complex systems, including transportation networks and Word Wide Web. Protein–protein interactions and metabolic pathways are also described as complex networks using graph modeling. Actually, these large graphs generally share common topological properties. General properties of big networks are summarized in the article of Mark Newmann, 2003 [2].

Graph-based analysis is an interdisciplinary approach; therefore, we may use the theory and methods, developed for graphs that have no obvious relevance to biology, medicine, or linguistics.

## Hypergraph

I have realized that concordance, obtained for current single, essentially represents the environment of corresponding hypergraph node. Concordance allows to examine the environment of current word within preliminary defined radius. Noteworthy, concordance shows the radius in the terms of letters, rather than hypergraph nodes. In contrast, Amorpha version 10 (V10) program allows to expand the nodes (not letters) adjacent to selected node of hypergraph. This idea will be explained in more details and illustrated later in this article.

<u>Thus, walk on reference singles from the top down allows to visit the nodes on linguistic hypergraph</u>.

The hypergraph is built via **<u>fusion</u>** of sentences by identical singles. Singles became the nodes, and the sentences became the edges of hypergraph. The fusion is performed for all sentences in source text together as they are, <u>without parsing</u>. An example of small linguistic hypergraph is shown on Picture 4.

**Picture 4.**
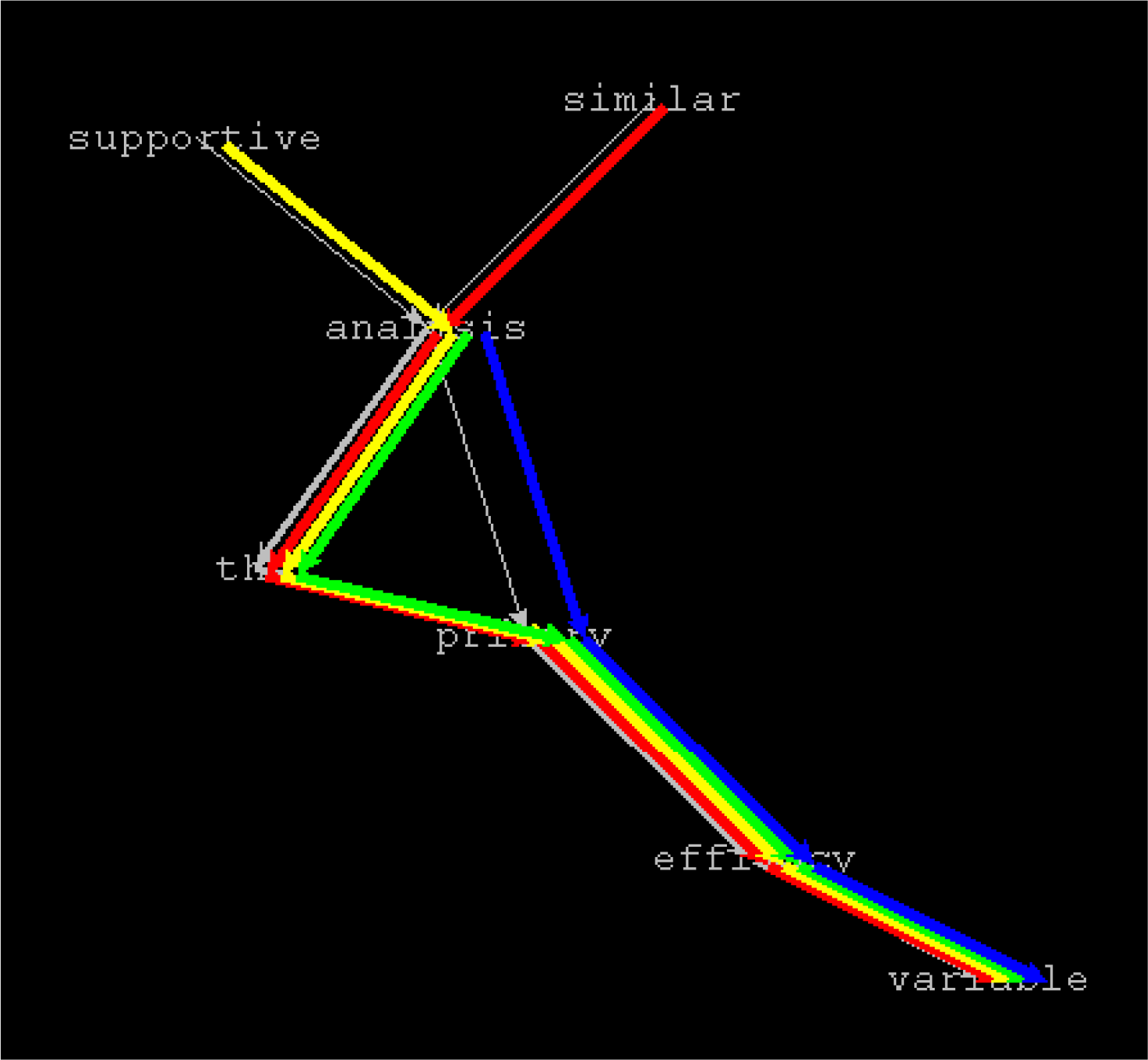
Linguistic hypergraph, produced in Amorpha version 8. The preposition “of” was excluded.

Preliminary parsing is avoided deliberately, because any syntactic or semantic parsing procedures inevitably fail in a number of cases. This is explained by the reasons, well known in linguistics: variation (more than one way of saying the same thing), ambiguity (same words with different meanings), as well as wide range of mistakes in meaning, logics, style, and specific knowledge areas. Even the texts within focused corpora contain wide range of rules that are not necessary to follow.

Therefore, fusion of original sentences by identical singles is an **<u>exact</u>** and **<u>unambiguous</u>** procedure that provides solid and reliable base for linguistic analysis.

Inversely sorted Refsi List allows to focus our attention on key nodes of the hypergraph. Thus, creation of hypergraph without parsing and walk on its keys <u>ensure**s**</u> control on a text. The concept of hypergraph is the key concept for analytical programs, included in Amorpha software package.

## The structure of hypergraph

By definition, hypergraph is a graph, in which one edge can include more than 2 nodes. Formally, a hypergraph H is a pair H = (X, E) where X is a set of nodes and E is a set of edges. According to another definition, hypergraph can be represented as a bipartite graph, in which the nodes from one part will correspond to the nodes of hypergraph, while the nodes from another part will correspond to the edges of hypergraph.

Therefore, according to definition of hypergraph as bipartite graph, each node of hypergraph is directly connected to all his edges. Each edge, in turn, is directly connected to all his nodes. Therefore, each node of hypergraph is directly connected to all nodes that belong to his edges. In other words, each node of hypergraph has the information about his <u>environment</u> just after building of hypergraph.

In linguistic hypergraph, created by fusion of all sentences, the nodes are singles (words and non-word elements) and the edges are sentences. In this hypergraph, each word is directly connected to its context: all words before and after current word in all sentences that have this word.

In contrast to hypergraph, in classic graph an edge always includes exactly 2 nodes. As the result, classic graph, in classic graph a node is directly connected only to its nearest neighbors. Therefore, extensive walk on classic graph is required to get the information about overall structure of the graph.

### Hypergraph and summary graph

Linguistic hypergraph can be analyzed as classic graph after conversion to **summary graph.** Summary graph is produced from hypergraph by fusion of all <u>edges</u>. If two adjacent nodes in hypergraph are connected with more than one edge, these edges are joined to produce one edge in resulting summary graph. Each edge of summary graph has weight: edge weight is directly proportional to the number of different edges between the two nodes in original hypergraph. For example, fusion of 5 different hypergraph edges between the two adjacent nodes will produce one edge in summary graph with weight equal to 5.

## Graph Visualization

Visualization is very important for analysis of large graph or hypergraph. A graph that has more than some few nodes is just impossible to imagine (see Picture 5 below).

**Picture 5.**
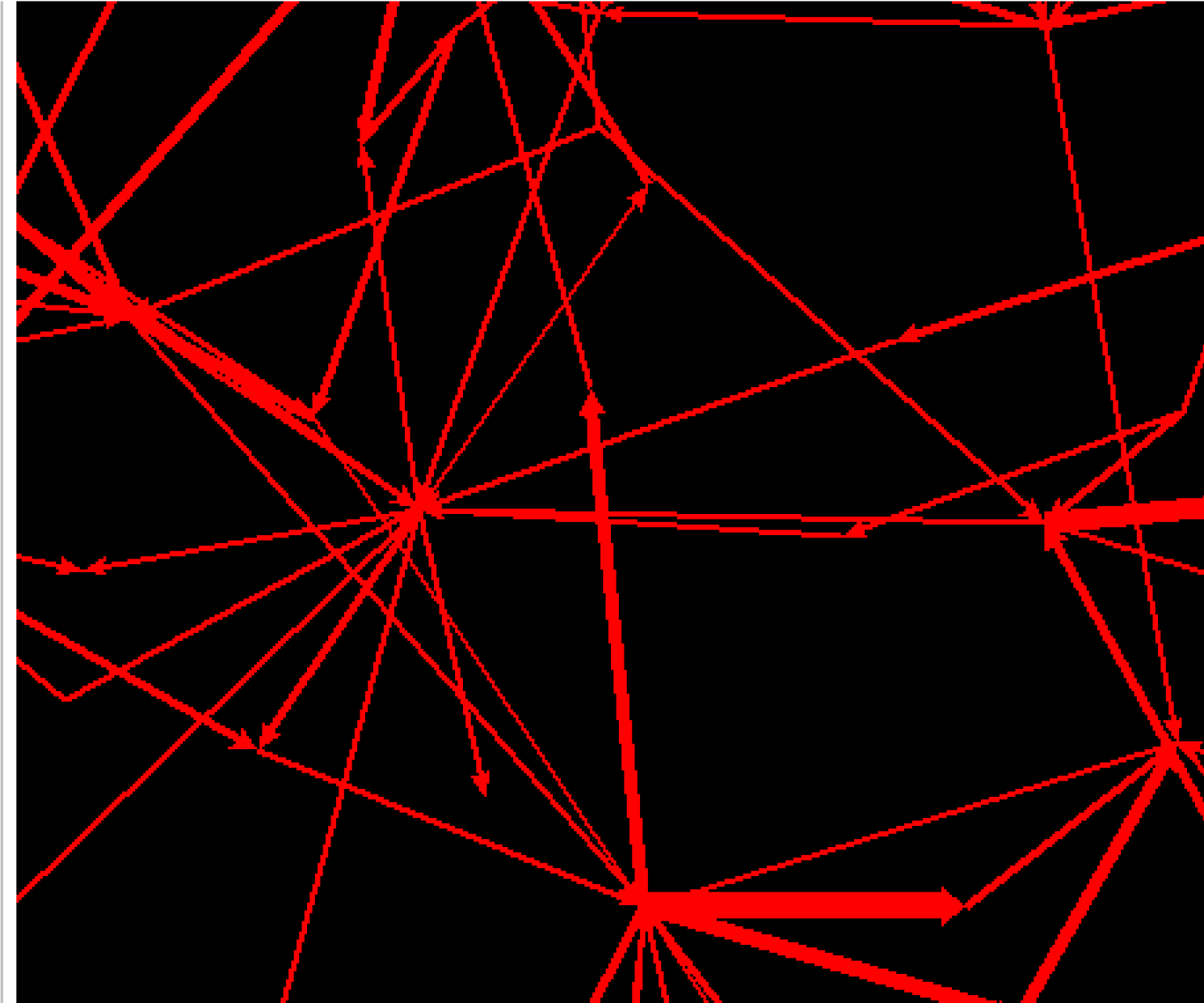
Core structure of summary graph formed by fusion of all sentences that contain the word “efficacy”. Words are not shown to demonstrate overall structure of the graph. The graph was built using Amorpha version 9 on PsAr corpus.

In fact, the articles, describing excellent algorithms for graph visualization, appeared in early 90-s (Kamada-Kawai [7], Fruchterman-Reingold [8]). However, many scientific articles and books about graphs still describe the graphs without any visualization or with poor, insufficient visualization.

The first algorithm for force-directed graph visualization was proposed by Japanese scientists Kamada and Kawai in 1989 [7]. This algorithm is essentially similar to the algorithms of geometry optimization, used in molecular modeling. In general, graph nodes are represented as particles with electric charge, while the edges are represented as springs. The algorithm of Kamada-Kawai was further improved by Fruchterman and Reingold [8]. Fruchterman-Reingold method has excellent implementation in Igraph software package for graph analysis (Gabor Csardi, Tamaz Nepusch, 2010 [5]).

Geometrical layout for summary graph can be created layout using Fruchterman-Reingold (FR) algorithm. Then the summary graph can be visualized as the whole graph, or as parts of this graphs. In large graphs, visualization of complete graph can be impractical or even impossible when total number of nodes or edges is too high. In this case, visualization of some relevant parts of the graphs (<u>subgraphs</u>) is the optimal method.

## Small-World Graphs

The shape of large linguistic summary graph was described in the article of Jean Veronis, 2007 [3]. This article makes the bridge between the two different worlds: linguistics and network theory.J. Veronis mentioned the pivotal article of Mark Newmann, 2003 [3] that describes common properties of so-called “small-world” graphs. Small-world graphs were previously defined in the articles of Barabási & Albert, 1999 [9] and Watts & Strogatz, 1998 [10]. Small-world graphs model were known to describe Word Wide Web, transportation networks, social networks, and biological networks. However, this model was not previously discussed in the context of linguistics.

All small world graphs contain clusters. The nodes inside such clusters have many connections to each other (highly interconnected nodes). However, there are few connections between different clusters. In social networks, such clusters are formed by friends that know each other. In linguistic summary graph, the clusters are formed by words that closely related to each other. Usually, these words comprise phrases, corresponding to key concepts of a text. An example of the structure of small world graph is shown in Picture 6 below.

**Picture 6.**
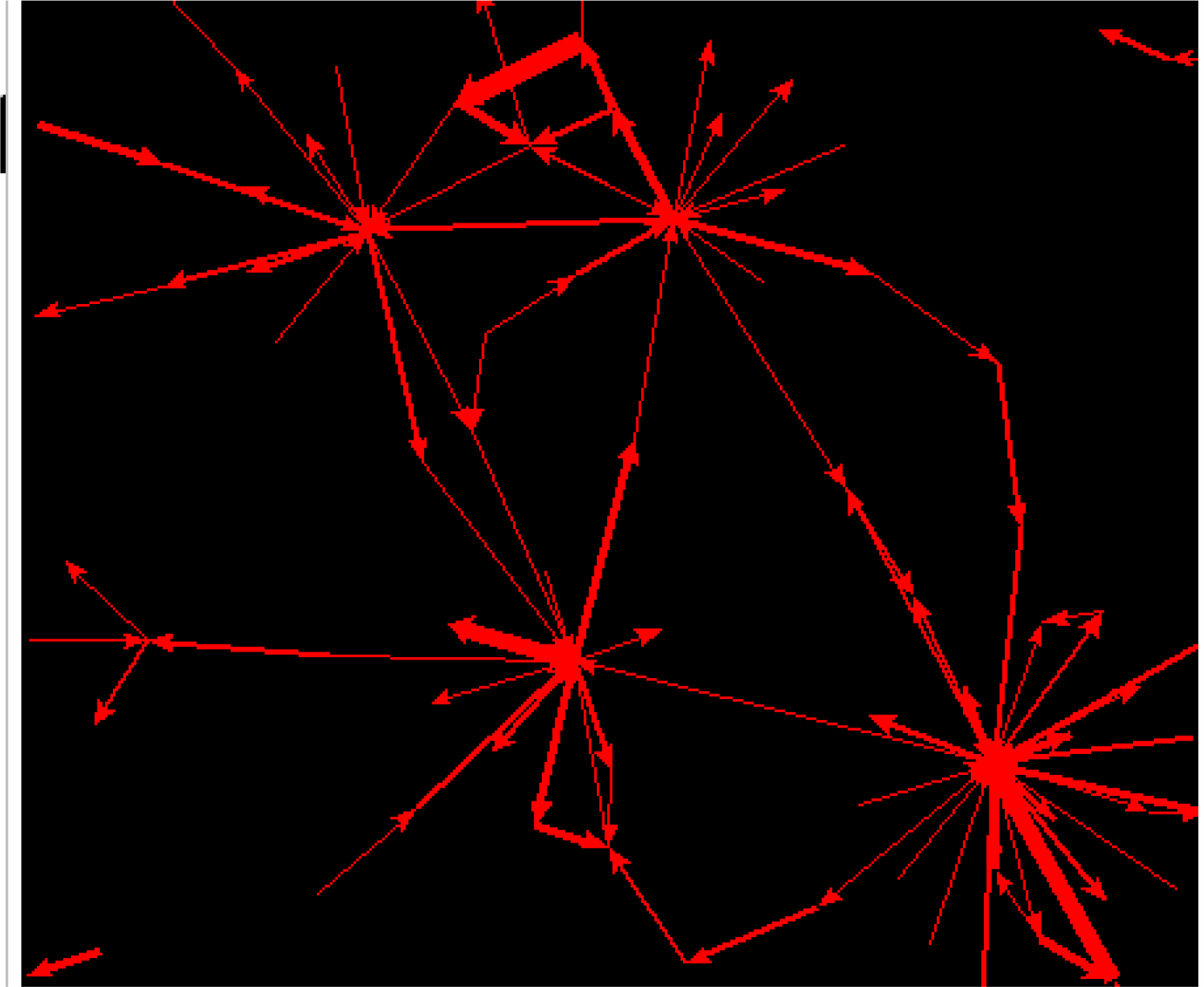
The core of linguistic summary graph demonstrates typical structure of small-world graph. The graph was built on PsAr Corpus in Amorpha version 9. To get this graph, minimal edge weight was limited to 42.

Jean Veronis performed deep analysis of mathematical aspects of graph theory that were discussed in the context of network theory. In his article, the concept of small-word linguistic graphs was used for word sense disambiguation (WSD). Jean Veronis demonstrated that once a text is represented as summary graph, this graph follows “small-world” criteria. Analysis of <u>clusters</u> of a big linguistic summary graph allows clearly identify the “islands” of closely related words.

In addition, Bales and Johnson [4] provided a review of on large-scale semantic networks, including real-world (not artificially created) networks. These authors showed that 15 of 28 (53.6%) original articles, included in the review, mentioned evidence of “small-world” characteristics of investigated networks. This review confirmed that networks, generated from natural language, share common topological properties with previously discussed transportation, social, and biological networks.

My work extends the ideas of Jean Veronis; this article is aimed to demonstrate that representation of a text as summary graph has more potential implementations beyond WSD. Summary graphs of big texts follow the small-word criteria; this is solid and reliable basis for exact analysis of a text or focused corpus. Therefore, the concept of linguistic analysis using small-world summary graphs is extended from WSD to full-scale linguistic analysis.

## METHODS

### Methods of linguistic analysis

In general, 4 different methods of linguistic analysis are discussed in this article:

- examination of summary graph based on <u>edge weight</u>
- examination of summary graph based on <u>distance</u> from selected node
- direct examination of <u>word frequency list</u> (Refsi List)
- search of relevant sentences in original articles using <u>sequential multiple filter</u> with target words or phrases

#### Edge weight as the criterion

This method is implemented in <u>Amorpha version 9</u> (V9) program. The method is based on slicing of summary graph according to edge weight. Initially, minimal edge weight threshold is specified. The resulting subgraph contains only edges with the weight above the threshold. If minimal edge weight was selected near the maximal edge weight in a whole graph (the most “heavy” edges), resulting subgraph contain only the most important terms or concepts in this graph. Gradually lowering minimal edge weight threshold, we can add another important terms to the subgraph. At some stage, the subgraph became too big for easy visual examination. If required information was not yet obtained, the program can produce a subgraph of current graph using additional filters on the range of sentences, comprising the graph. An example of core graph, produced in V9, is shown in Picture 7 below.

**Picture 7.**
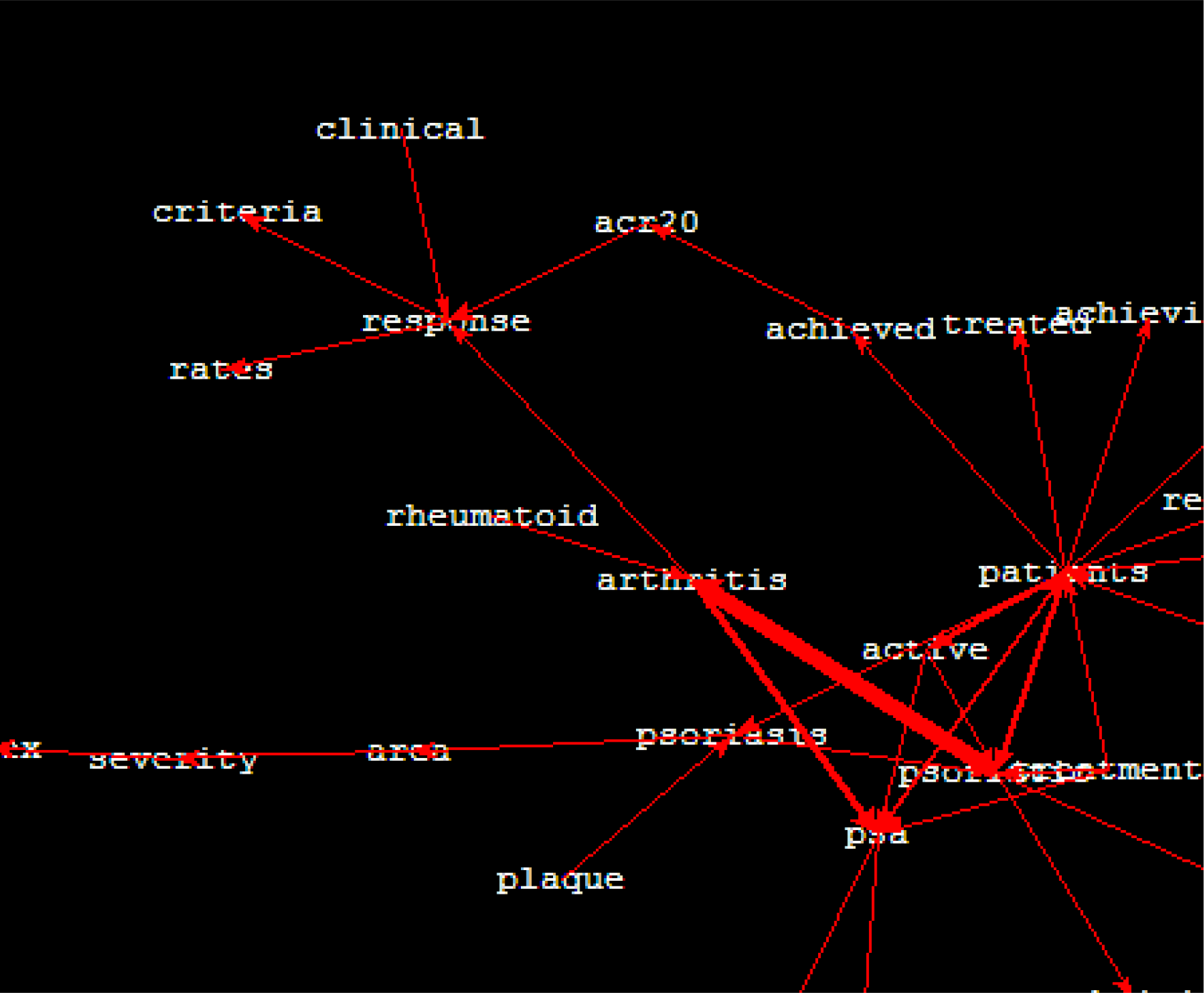
View of central core of PsAr Corpus in Amorpha version 9; all sentences included, minimal edge weight is 19.

#### Distance from selected node as the criterion

This method is implemented in <u>Amorpha version 10 (V10)</u> program: the essential strategy is to expand the context (environment) of selected node gradually, step by step, using the <u>distance</u> from selected node as the criterion. In V10, the distance if defined in the <u>number of nodes</u>, adjacent to selected node.

This methods allows rapidly identify all essential information, directly related to the term of interest. For example, in a corpus focused on specific drug the information about current spectrum of diseases treated with this drug can be immediately obtained by expansion of key word, related to disease, such as “**<u>cancer</u>**” (for anti-cancer drug corpus), or “**<u>lymphoma</u>**” (for drugs approved to treat some type of lymphoma), or may be more general terms (“disease”, “disorder”, etc.).

The results are provided as text view and graph view. Text representation includes a list of fragments that contain the context words, selected according to specified distance of expansion. This list of fragments is obtained using previously described concept of hypergraph. The sentences represent the edges of hypergraph; the algorithm of V10 just captures the words that lying on the left side and/or on right side from selected node within specified distance. The distance of expansion is interactively regulated using two scales (Left Passage Scale and Right Passage Scale). Representation of text in V10 as concordance and graph is shown in Picture 8 and Picture 9, respectively.

**Picture 8.**
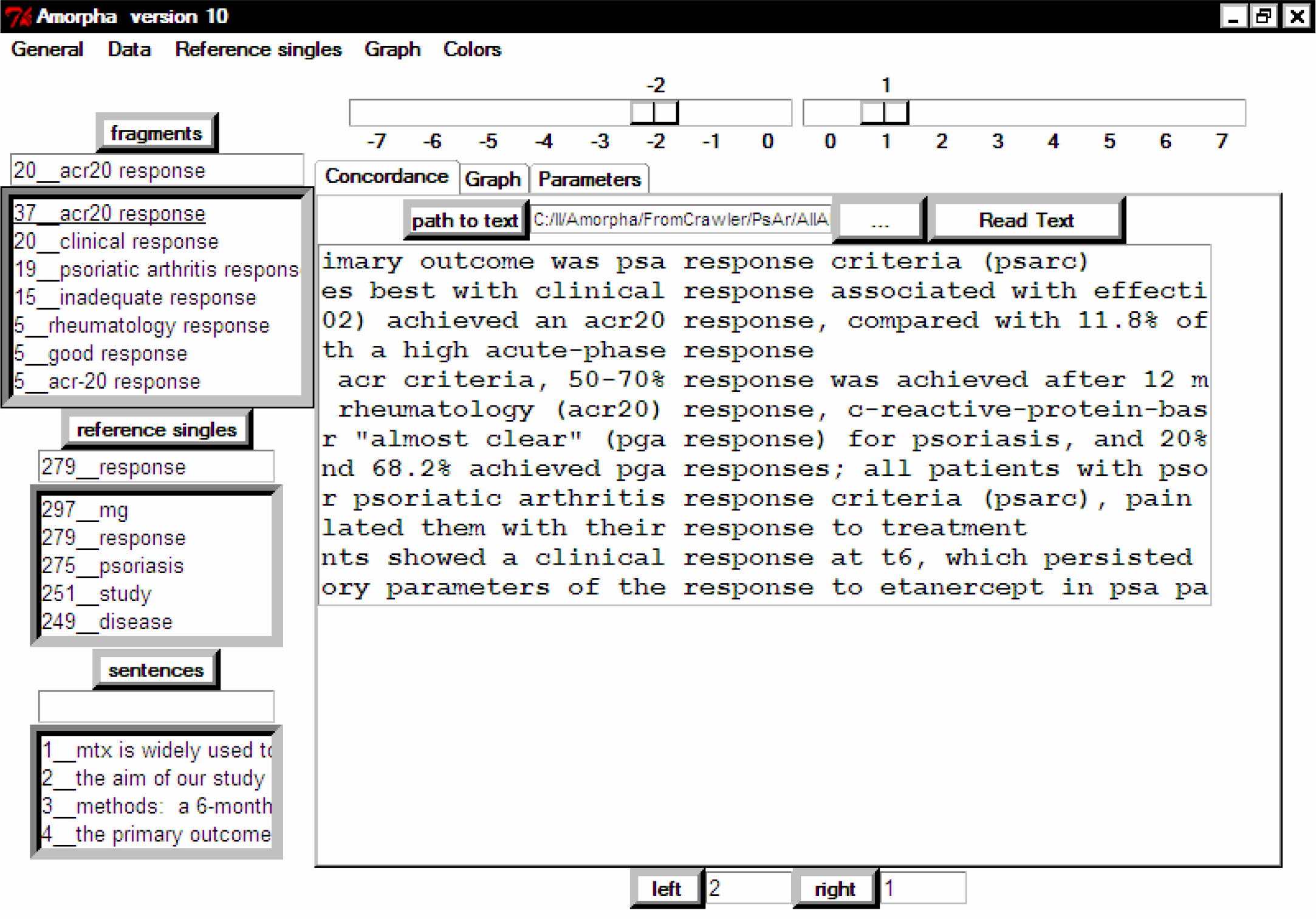
Expansion of “response” single in V10: concordance view. In this picture the expansion distance is specified as 2 steps left and 1 step right from the center. First step right will capture the central single (“response” in this case). Zero steps right means that central single is not included to the results. This can be useful when the resulting graph is too big; in this case, elimination of central single will reduce density of the graph and facilitate visual examination. The fragments, obtained for specified expansion distance, are shown in “<u>fragments</u>” window. Current single is selected in “reference singles” window.

**Picture 9.**
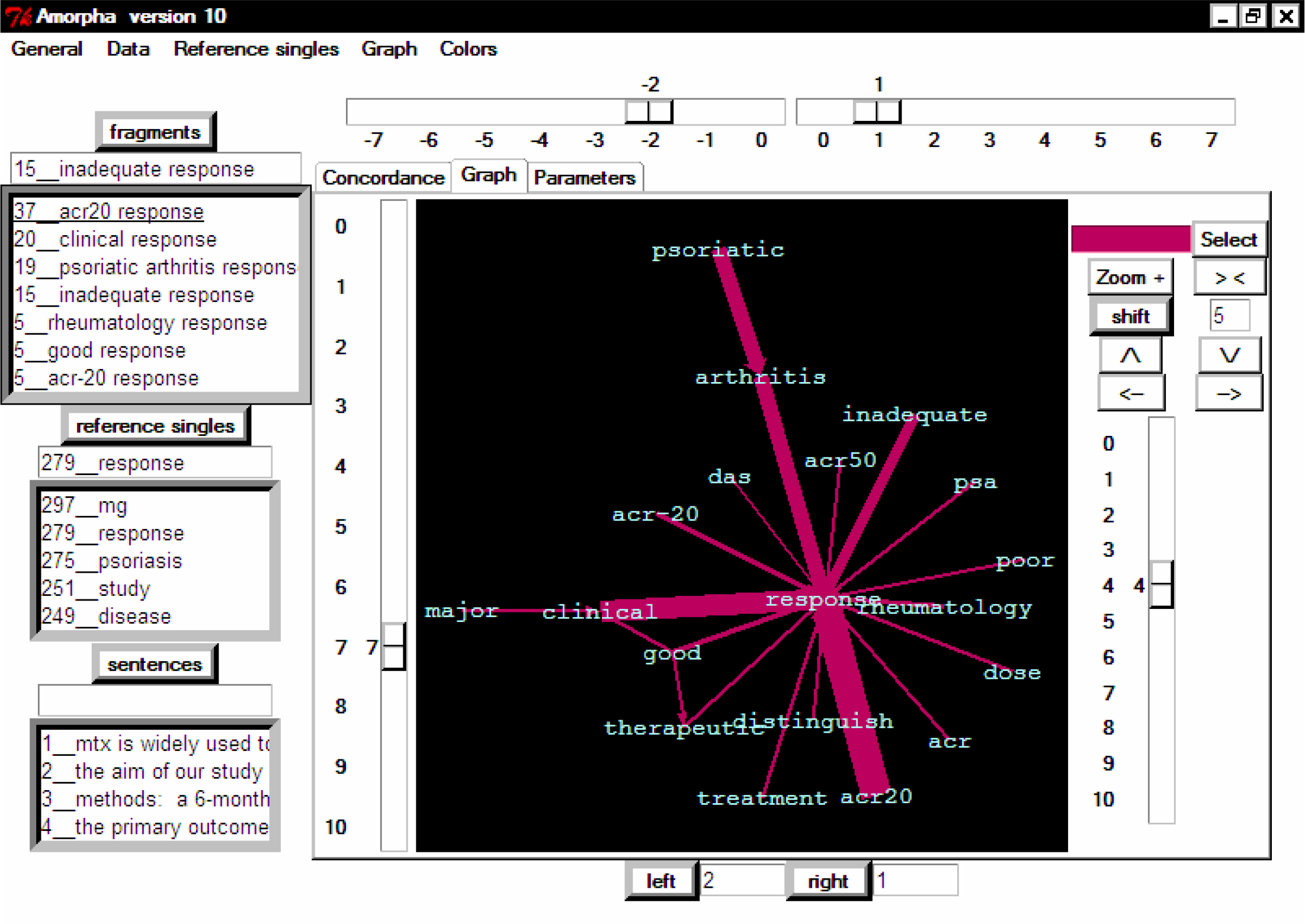
Expansion of “response” single in V10: graph cluster, associated with response. Two steps left and one step right were taken. Graph representation is equivalent to fragments collection, showed on the image above. To make the graph easy to understand, minimal edge weight was set to 3; therefore, all fragments with frequency 2 or 1 were not included in this graph. Edge weights are proportional to the frequency of initial fragments.

In this example, two steps left and one step right were taken. Resulting text fragments are extracted and sorted by frequency (“fragments” window on the top). The results show that ACR20 is the main criterion of response for psoriatic arthritis.

This analysis also provides additional terms that can be used in subsequent searches, such as “inadequate response” and “rheumatology response”.

Left Passage Scale and Right Passage Scale are located above the concordance view. Each change in these scales will immediately produce new collection of fragments, providing easy and rapid navigation on the text.

Noteworthy, concordance view includes all words and symbols that really present in the text. However, in “fragments” window the resulting fragments will not include prepositions, conjunctions, articles and modal verbs. As mentioned previously in this article, these high-frequency singles visually attract attention on the graph and increase total amount of different fragments. Therefore, elimination of these singles facilitates and accelerates interpretation of the results.

Another example is a corpus, focused on specific disease. In such corpus, V10 allows rapidly identify current therapeutic strategies to treat the disease. This can be done by expansion of key word, directly related to drug, such as dose measurement unit, for example, “**<u>mg</u>**”. The resulting subgraph of nearest environment will show the list of drugs that were actually used for treatment of the disease, with indication of therapeutic doses. Further extension to the right side will provide the information about treatment regimen, such as, for example, “once a day”, “twice a week”, etc. Relevant results are presented later in this article.

#### Direct examination of word frequency list

This method is implemented in all versions of Amorpha software. For example, we can use this method to find top drugs or all drugs, mentioned in a text. The words can be compared with already existing list of drugs. Alternatively, the words can be searched for drug-specific suffixes, such as “ine”, “ole”, etc. For example, using this approach, we can easily identify the drugs, based on monoclonal antibodies, using specific “mab” suffix to search drug names.

#### Search of target phrases in all articles

All relevant sentences can be found using local search with previously identified key terms; this method is implemented in <u>Amorpha version 11</u> (V11) program.

In contrast to V9 and V10, where collection of texts is analyzed as one big text, V11 keep information about the source articles, including authors, titles, and PubMed indices. Local search of relevant sentences in V11 can be further refined using filters with relevant words of phrases, sequentially applied to select sentences with highest relevance to specified query.

### Software

Amorpha software was written by the author of this article in the period from 2010 to 2017. This software was written in pure Python. Igraph software package [5] is the only external library that was integrated to Amorpha software. All the remaining components of Amorpha were written using standard builtin functions of Python 2.6.6. The software runs on Windows and have graphic user interface, based on Tkinter [6] (included in standard Python library). Graph visualization procedures are also written entirely in Tkinter.

### Source texts

Source texts were retrieved from PubMed as abstracts of scientific articles on clinical trials. PubMed search terms are listed in the table below:

| Search aim | Search terms | Name of collection of texts for analysis | Total number of articles in PubMed | Number of articles, taken for analysis |
| --- | --- | --- | --- | --- |
| psoriatic arthritis | "psoriatic arthritis"[Title]<br>AND Clinical Trial[ptyp] | PsAr<br>Corpus | 131 | 131 |
| Therapy of breast cancer | "breast cancer"[Title] AND ("therapy"[Title] OR treatment["Title"])<br>AND Clinical | Breast Cancer<br>Corpus | 2773 | first* 400 articles |
|  | Trial[ptyp] |  |  |  |
| paclitaxel | "paclitaxel"[Title]<br>AND Clinical<br>Trial[ptyp] | Paclitaxel<br>Corpus | 3274 | first* 400<br>articles |

\*first corresponds to the most recent articles in the list of articles, sorted by publication date

The abstracts were obtained using PubMed-specific focused web crawler that was designed to get only limited amount of abstracts from PubMed, according to initially defined PubMed search. This web crawler is a part of Amorpha software. As well, the other functional modules of Amorpha, it is small program, written in pure Python.

### Extraction of essential information

The following essential information was extracted:

- drugs used for treatment of breast cancer
- drugs used for treatment of psoriatic arthritis (PsAr)
- diseases treated with paclitaxel
- main risk factors, associated with breast cancer

The walk on summary graph, using the <u>distance from selected node</u> as the criterion, was the main method, used to extract this information. Examination of word frequency list was used to identify drugs, related to breast cancer.

## RESULTS

### Drugs for treatment of psoriatic arthritis

The results were obtained in V10 using expansion of 2 steps left and 1 step right from “<u>mg</u>” single. The drugs, used for treatment of psoriatic arthritis, with actual therapeutic doses, are shown in the table [Table 2] and graph [Picture 10] representations below.

**Table 2.** Drugs used for treatment of psoriatic arthritis

| **Drug name and dose** | **Frequency** |
| --- | --- |
| golimumab 50 mg | 15 |
| etanercept 50 mg | 13 |
| apremilast 20 mg | 11 |
| secukinumab 300 mg | 8 |
| adalimumab 40 mg | 7 |
| ustekinumab 45 mg | 6 |
| celecoxib 400 mg | 6 |
| etn 50 mg | 4 |
| etanercept 25 mg | 4 |
| brodalumab 140 mg | 4 |
| czp 400 mg | 2 |
| czp 200 mg | 2 |
| clazakizumab 25 mg | 2 |
| secukinumab 150 mg | 1 |
| onercept 50 mg | 1 |
| nim 200 mg | 1 |
| nim 100 mg | 1 |
| golimumab 50/100 mg | 1 |
| golimumab 100 mg | 1 |
| clazakizumab 100 mg | 1 |
| celecoxib 200 mg | 1 |
| brodalumab 280 mg | 1 |
| auranofin 6 mg | 1 |

**Picture 10.**
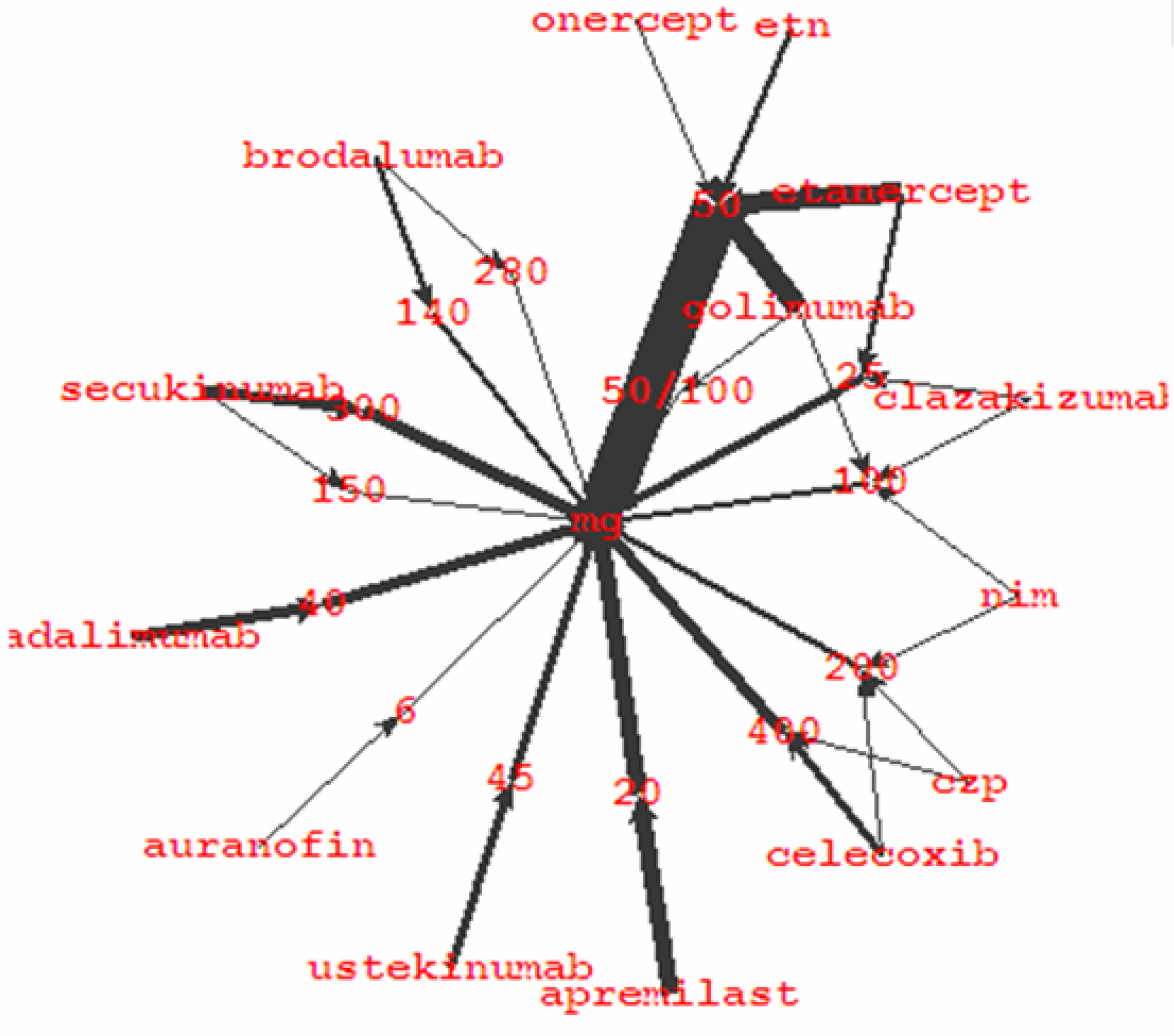
Drugs used for treatment of psoriatic arthritis. Graph image was produced in Amorpha version 10.

These results showed that 50 mg is the most frequent therapeutic dose for treatment of PsAr; etanercept and golimumab are the most frequently used drugs for treatment of PsAr.

The resulting graph is even more representative that table, because the graph shows different drugs used in the same doses, with summation of contribution of these doses to treatment trends. For example, onercept, etanercept, and golimumab are used in the same 50 mg dose.

### Drugs for treatment of breast cancer

The results were obtained using direct examination of word frequency list in V10. Top drugs, used for treatment of breast cancer, are shown in Table 3. The numbers of different articles for each drug were calculated in V11.

**Table 3.**
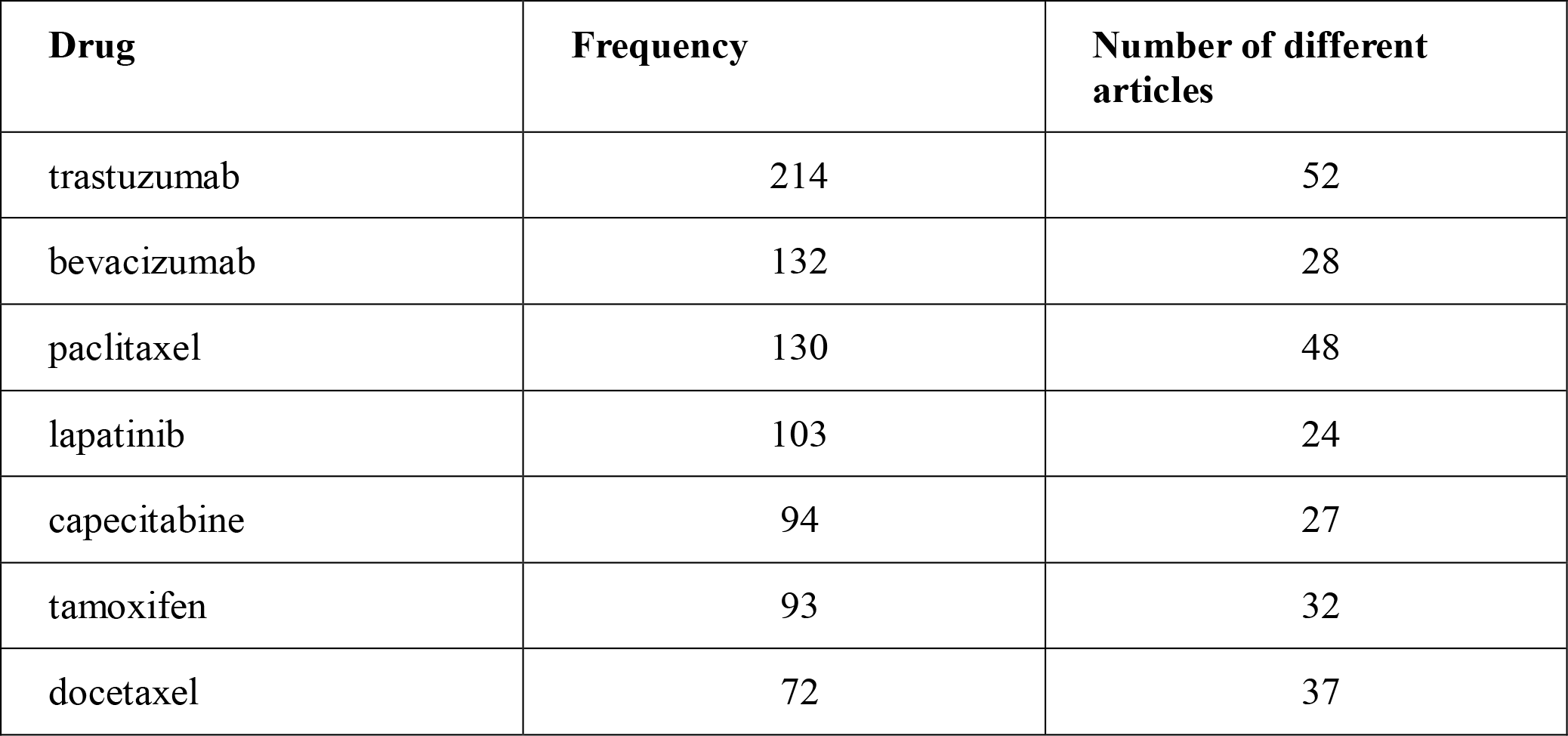
Drugs used for treatment of breast cancer

| **Drug** | **Frequency** | **Number of different articles** |
| --- | --- | --- |
| trastuzumab | 214 | 52 |
| bevacizumab | 132 | 28 |
| paclitaxel | 130 | 48 |
| lapatinib | 103 | 24 |
| capecitabine | 94 | 27 |
| tamoxifen | 93 | 32 |
| docetaxel | 72 | 37 |

Within selected set of articles (Breast Cancer Corpus), trastuzumab is the leading drug for treatment of breast cancer. Overall frequency of the word “trastuzumab” in Breast Cancer Corpus is 214; trastuzumab is mentioned in **<u>52</u>** different articles.

Paclitaxel has similar word frequency to bevacizumab (130 vs. 132); however, paclitaxel is mentioned in **<u>48</u>** different articles as compared with only 28 for bevacizumab.

### Paclitaxel

The spectrum of diseases, treated by paclitaxel, is presented in Table 4 below and corresponding graph [Picture 11]. The results were obtained in V10 using extension 1 step left and 1 step right from “cancer” single.

**Table 4.** Spectrum of diseases treated with paclitaxel

| **Disease** | **Frequency** |
| --- | --- |
| breast cancer | 143 |
| lung cancer | 78 |
| gastric cancer | 51 |
| ovarian cancer | 39 |
| cervical cancer | 37 |
| pancreatic cancer | 18 |
| endometrial cancer | 11 |
| urothelial cancer | 7 |
| peritoneal cancer | 3 |
| oesophageal cancer | 3 |
| gynecologic cancer | 3 |
| bladder cancer | 2 |
| thyroid cancer | 1 |
| stomach cancer | 1 |
| oropharyngeal cancer | 1 |

**Picture 11.**
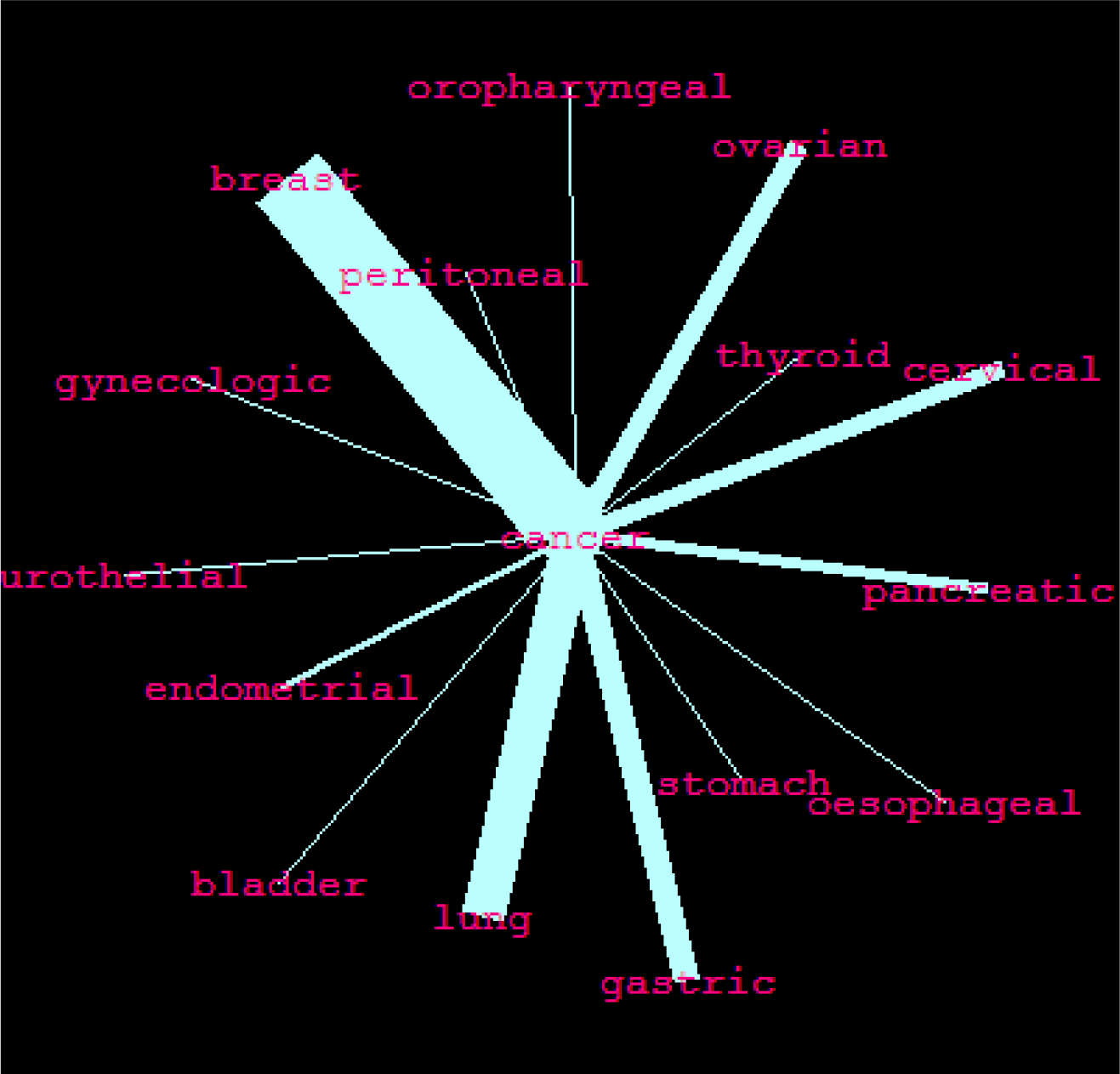
Spectrum of diseases treated with paclitaxel. Graph image was made in Amorpha version 10.

These results indicate that paclitaxel is most frequently used for treatment of breast cancer, lung cancer, gastric cancer, ovarian cancer, and cervical cancer. On the other hand, paclitaxel is rarely used to treat other types of cancer, such as thyroid cancer.

#### Risk factors of breast cancer

To get relevant information about risk, associated with breast cancer, all sentences with the word “risk” were extracted in V11 to form a new text. Then this text was loaded to V10; reference singles list was produced and analyzed. The word “estrogen” has highest frequency in the list among potential risk factors. To evaluate this hypothesis, all 400 articles of Breast Cancer Corpus were examined in V11. The word “risk” was used as the first filter to select all sentences with this word. These “risk” sentences were selected from 67 articles to form “all__risk” set of sentences. The next filter was applied to “all__risk” set using the word “estrogen” as second filter word. This new filter produced only 9 sentences that contain both “risk” and “estrogen”. These sentences were extracted from 4 different articles, and all sentences (except one sentence), actually discussed the risk of breast cancer related to estrogen. Pubmed indices of relevant articles are presented in Picture 12.

**Picture 12.**
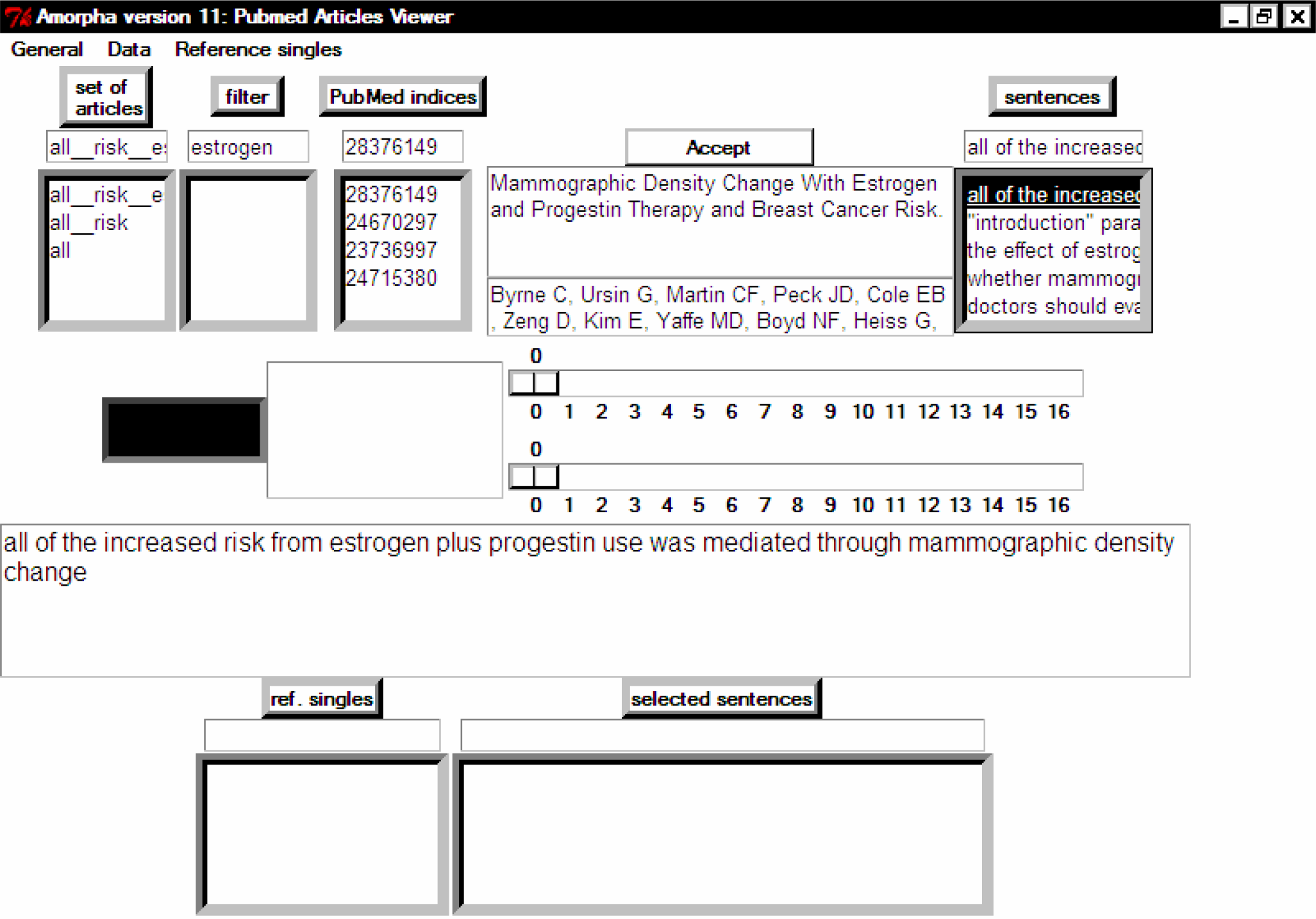
Extraction of sentences discussing the risk associated with estrogen in Amorpha version 11

#### Discussion

This article describes two methods used to explore linguistic summary graph:

1. examination of summary graph using edge weight as the criterion
2. examination of summary graph using the distance from selected node as the criterion

In fact, second (distance-based) method appears to be more useful, when the aim is to extract specific information from a text. However, the first method provides panoramic view of the whole text or big subgraph, and allows immediately identify the most important terms of a text. These key terms may be obvious for a scientist who perfectly knows this area of knowledge. However, for learning of new knowledge area, this immediate visualization of key concepts is very useful. This method will be helpful for scientists as well as for medical writers, starting to explore new knowledge area. For example, microbiologist can use this method to get key terms in oncology or neurology.

### Breast cancer

Based on distribution of top drugs for treatment of breast cancer, additional search in PubMed was performed for clinical trials, conducted with <u>paclitaxel</u> (the second most frequent drug for treatment of breast cancer). This is an example of validated search refinement, based on linguistic analysis of an initial text.

### PsAr

Focused subgraph of summary graph, presented in this article, demonstrates current trends in therapy of PsAr. Once a summary graph was created, less than a minute (actually a few seconds) is required to get these results. Noteworthy, that original collection of texts, PsAr Corpus, contains 131 abstracts. In the absence of Amorpha software, the analysis of such amount of abstracts will require significant time and efforts.

### Paclitaxel

Paclitaxel was included after examination of the list of drugs, used to treat breast cancer, because paclitaxel is the second most frequently used drug for treatment of breast cancer.

Rapid quantitative assessment of the disease spectrum for specific drug allows to understand strength and weakness of this drug. Noteworthy, the diseases in the “tail” of fragment distribution curve may be even more interesting that the diseases in the peak, because “tail” (low-frequency) diseases can represent an unmet medical need. This hypothesis should be further evaluated using additional refined literature search and analysis of selected articles.

If the initial hypothesis about unmet medical need for a disease is confirmed, this information can indicate a perspective direction of new drug discovery or clinical trials for already existing multi target drug candidates.

### Risk of breast cancer

V9 and V10 provide key terms, directly relevant to the area of interest. This allows to refine the search criteria and, finally, to improve results of the search for scientific literature. In contrast to V9 and V10, where collection of texts is analyzed as one big text, V11 keep information about the source articles, with authors, titles, and PubMed indices. Importantly, local search of relevant sentences in V11 can be further refined applying relevant words of phrases sequentially. Application of sequential filters in V11 takes a few seconds and produces highly relevant sentences that can be subsequently used for writing scientific summary or report.

### Conclusion

Linguistic analysis of scientific articles and regulatory documents using Amorpha software provides exact, quantitative results, including evaluation of intrinsic trends. This information can be used for making strategic decisions on drug development.

Rapid and exact analysis of current trends allows to save money and efforts, and focus the process of drug development on the most promising issues.

## Footnotes

If you are interested in new analysis with Amorpha software, please contact the author using the.

